# Optimized Workflow for Self-Delivering siRNA-Mediated Gene Knockdown in Unstimulated Human CD4+ T Cells

**DOI:** 10.1101/2025.07.11.664322

**Authors:** Balasiddaiah Anangi

## Abstract

T-cell-based therapies have transformed cancer treatment, yet dissecting the molecular networks that govern T-cell function remains technically challenging. Gene manipulation in primary human T cells typically relies on electroporation or viral vectors, approaches that often compromise viability, alter cell physiology, or introduce genotoxic risks. Accell™ small interfering RNAs (siRNAs) are chemically modified, self-delivering siRNAs that bypass the need for transfection reagents or viral systems. Despite their potential, Accell siRNAs have been underutilized in human T cells. Here, using freshly isolated, unstimulated primary CD4+ T cells, we optimized conditions for Accell siRNA uptake and gene expression knockdown. We demonstrate efficient and reproducible delivery of Accell siRNAs across multiple serum-free media, with Accell delivery medium providing the highest uptake. Using a GAPDH-targeting Accell siRNA pool, we achieved robust transcript knockdown without loss of viability, and knockdown efficiency was modestly enhanced by CD3 stimulation. This streamlined workflow enables rapid assessment of gene function in resting human T cells without genetic modification or pre-activation, expanding the toolkit available for immunology and therapeutic research.

## Introduction

T cells are central regulators of immune responses, mediating protection against infection and cancer but also contributing to autoimmunity when dysregulated (1). Advances in T-cell engineering, including chimeric antigen receptor (CAR) and T-cell receptor (TCR) therapies, have revolutionized cancer treatment (1). However, further progress requires efficient, reproducible, and minimally disruptive methods to manipulate gene expression in primary T cells.

Two major technologies dominate gene function studies in T cells: CRISPR-Cas9–mediated knockout and small interfering RNA (siRNA)–mediated knockdown (2,3). Both approaches typically rely on electroporation or viral vectors, which pose significant challenges. Electroporation can reduce cell viability and requires prior activation of T cells, limiting its utility for studying resting cells (2,3). Viral vectors raise safety concerns, including insertional mutagenesis, and often involve labor-intensive production. Consequently, there remains a need for alternative delivery strategies that preserve cell health while enabling efficient gene silencing.

Accell™ siRNAs (Horizon Discovery) are synthetic, chemically modified siRNAs designed for self-delivery (4). Unlike conventional siRNAs, which require lipid- or polymer-based reagents, Accell siRNAs enter cells directly and are stabilized intracellularly, making them effective in hard-to-transfect cell types such as neurons and lymphocytes. To illustrate the difference between conventional siRNAs and chemically modified Accell siRNAs, a schematic overview is shown in Figure 1. Previous studies have demonstrated their utility in suspension and immune cells, including leukemic cell models (14), as well as in more complex 3D culture systems (15). Despite being available for nearly two decades, Accell siRNAs have seen limited use in primary human T cells. A small number of studies have applied them in T-cell research (5–13), often in pre-activated or expanded cells, leaving their performance in freshly isolated resting T cells underexplored.

**Figure 1.**
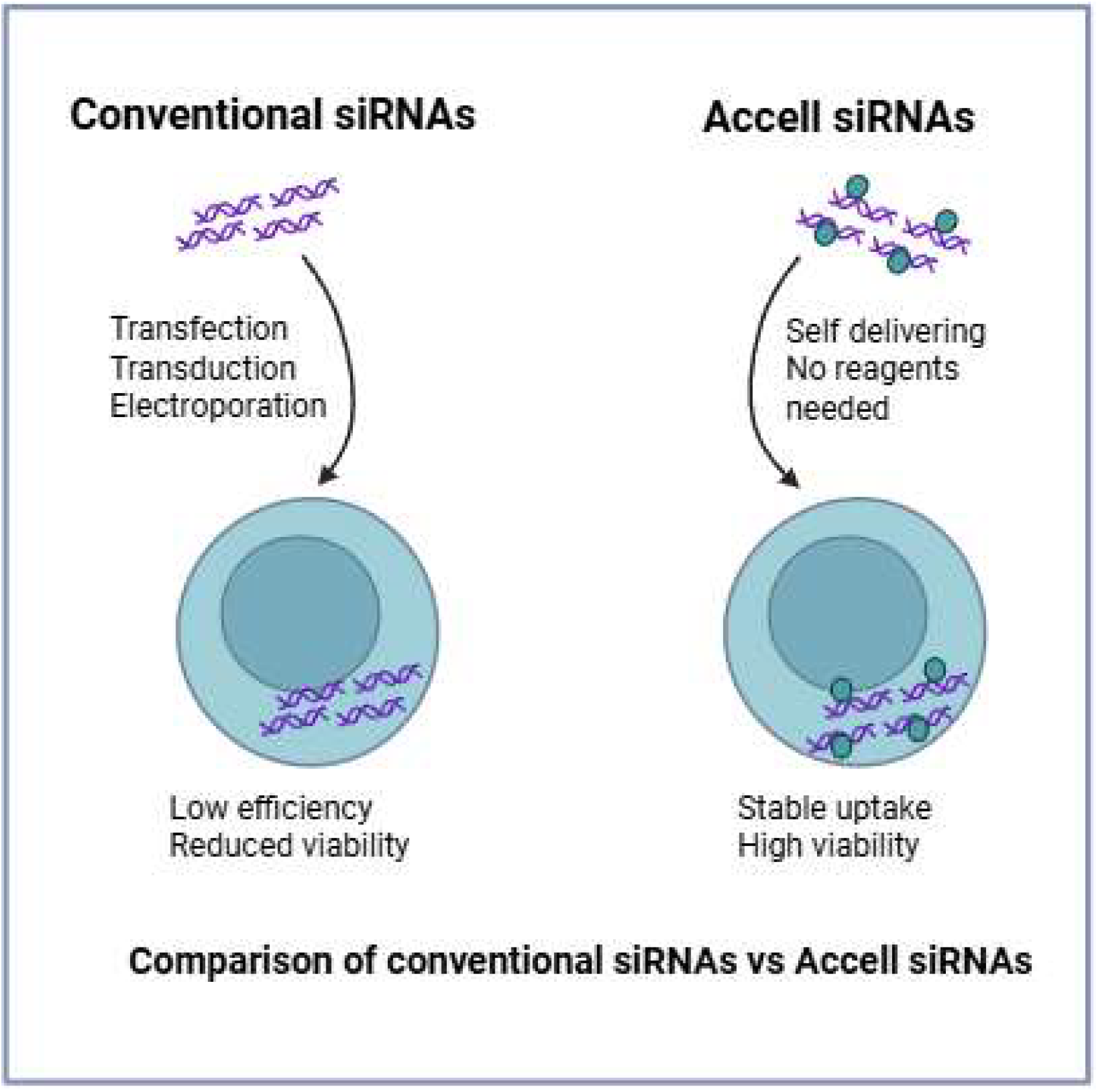
Comparison of conventional siRNAs and Accell siRNAs. Conventional siRNAs require delivery methods such as transfection, transduction, or electroporation, which often result in low delivery efficiency and reduced T cell viability. In contrast, Accell siRNAs are chemically modified for self-delivery, requiring no additional reagents, and enable stable uptake with high T cell viability.

Here, we systematically optimized Accell siRNA uptake and transcript-level knockdown in human primary CD4+ T cells without prior activation. We evaluated siRNA delivery across different serum-free media, quantified uptake kinetics, and validated gene knockdown efficiency using GAPDH as a model target. The optimized workflow provides a practical and reproducible approach for RNA interference in unstimulated human T cells, enabling rapid interrogation of gene function in settings directly relevant to immunology and therapeutic development.

## Results

### Accell siRNAs are efficiently delivered into unstimulated primary human CD4+ T cells

According to Dharmacon™, Accell siRNA delivery medium maintains cell health and provides suitable serum-free conditions for efficient uptake of Accell siRNAs. Although this medium is recommended for use with Accell siRNAs, Dharmacon also advises testing other serum-free media that are routinely used for the specific cell type of interest. Guided by this recommendation, we compared two widely used serum-free T cell culture media—X-VIVO™ 15 and TexMACS™— alongside the Accell siRNA delivery medium.

Freshly isolated CD4^+^ T cells were treated with Accell™ green non-targeting siRNA, a fluorescently labeled control siRNA that does not hybridize to any human transcript, to assess uptake efficiency. Cells were cultured in X-VIVO™ 15, TexMACS™, or Accell siRNA delivery medium, with or without CD3 stimulation. Uptake and viability were analyzed by flow cytometry at 24 h and 48 h. Accell siRNAs were robustly internalized under all tested conditions, with >95% cell viability maintained. Of note, the addition of siRNA did not affect cell viability compared to untreated cells, indicating that Accell siRNAs are not toxic to CD4^+^ T cells (Fig. 2a, Fig. 3a). Accell delivery medium produced approximately a threefold increase in mean fluorescence intensity compared to T cell media and resulted in more homogeneous uptake across the cell population (Fig. 2b, Fig. 3b). Anti-CD3 stimulation had no significant impact on siRNA delivery or cell viability. Uptake plateaued at 24 h and remained stable through 48 h, indicating that internalization is largely complete within one day (Fig. 2b; Fig. 3b). Together, these findings demonstrate that Accell siRNAs are efficiently delivered into primary human CD4^+^ T cells without compromising viability, and that Accell delivery medium provides optimal conditions for siRNA uptake.

**Figure 2.**
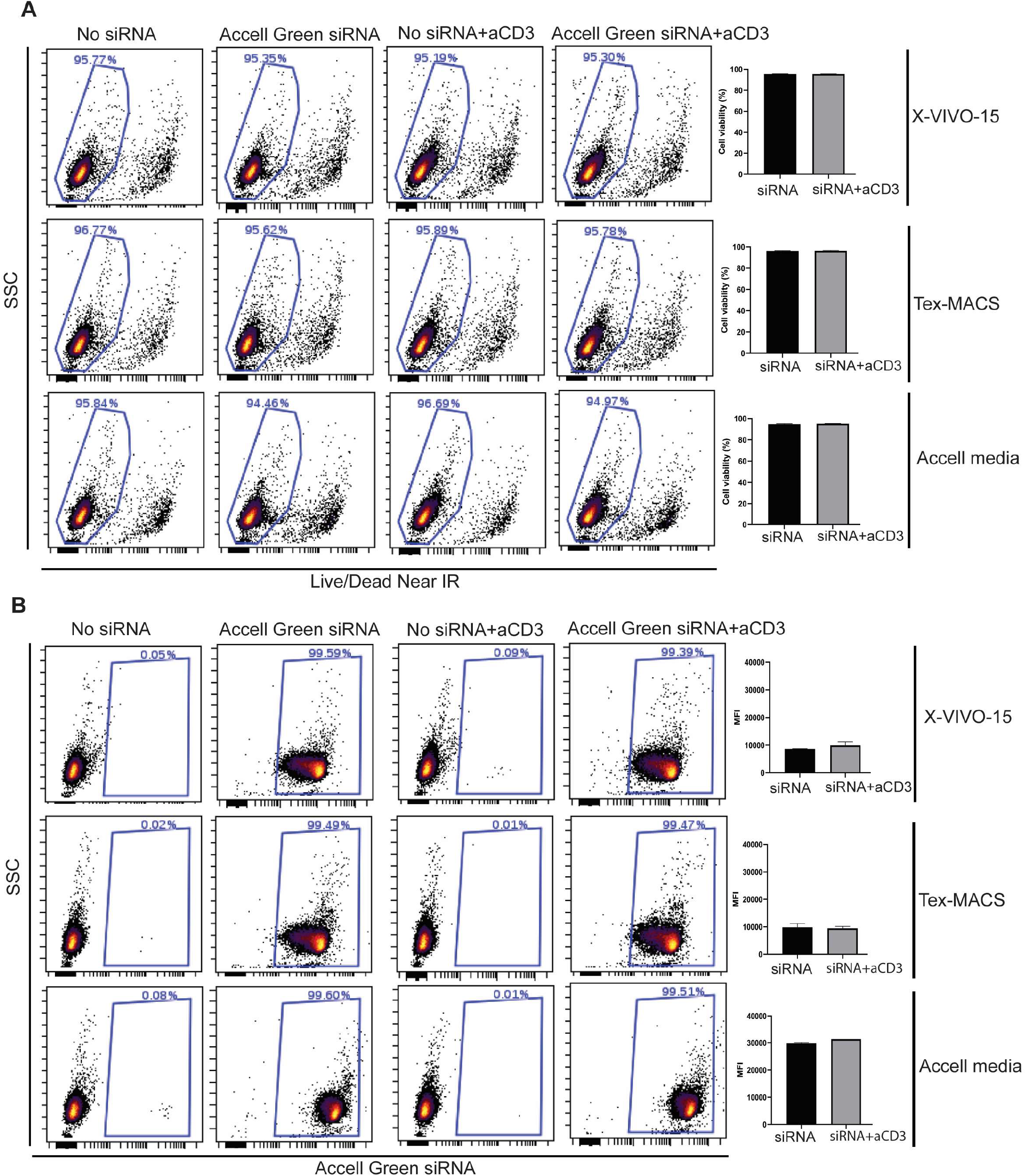
Flow cytometry analysis of purified human CD4^+^ T cells after 24 h treatment with Accell green non-targeting siRNA. Purified CD4^+^ T cells were cultured for 24 h in three different serum-free media (X-VIVO 15, TexMACS, or Accell medium) supplemented with or without αCD3 antibody. (A) T cell viability was assessed using Live/Dead Near-IR staining. Cells shown in the gates represent viable cells. Plots were gated on the total CD4^+^ T cell population. Quantification of T cell viability is presented in the graphs on the right. (B) siRNA uptake was measured as the percentage of T cells retaining the fluorescent siRNA signal. Mean fluorescence intensity (MFI) of the siRNA signal is shown in the graphs on the right. Plots were gated on the live T cell population shown in panel A.

**Figure 3.**
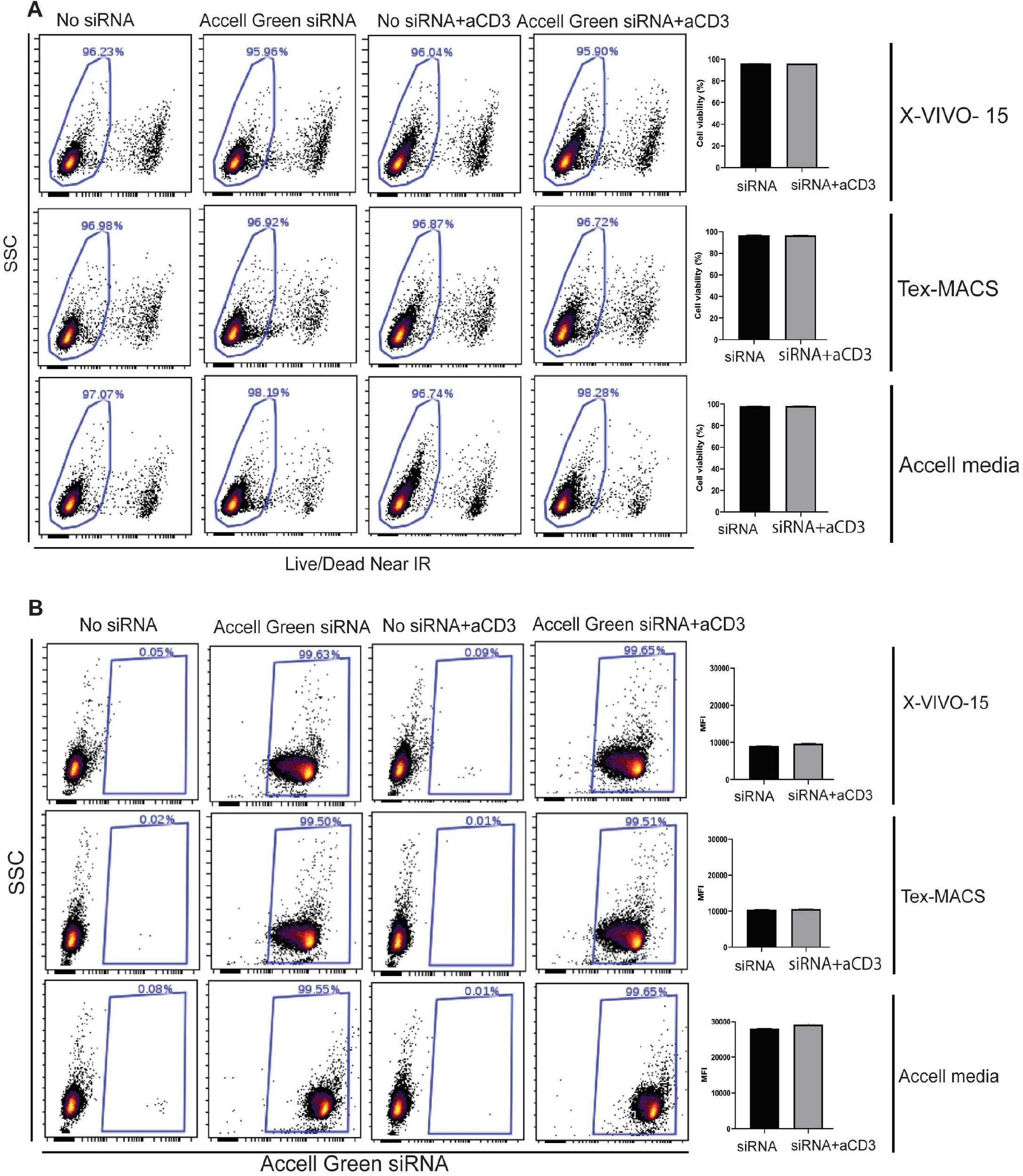
Flow cytometry analysis of purified human CD4^+^ T cells after 48 h treatment with Accell green non-targeting siRNA. Purified CD4^+^ T cells were cultured for 48 h in three different serum-free media (X-VIVO 15, TexMACS, or Accell medium) supplemented with or without αCD3 antibody. (A) T cell viability was assessed using Live/Dead Near-IR staining. Cells shown in the gates represent viable cells. Plots were gated on the total CD4^+^ T cell population. Quantification of T cell viability is presented in the graphs on the right. (B) siRNA uptake was measured as the percentage of T cells retaining the fluorescent siRNA signal. Mean fluorescence intensity (MFI) of the siRNA signal is shown in the graphs on the right. Plots were gated on the live T cell population shown in panel A.

### Accell siRNAs mediate efficient transcript knockdown in primary human CD4+ T cells

To test functional silencing, CD4+ T cells were treated with a GAPDH-targeting Accell siRNA pool or non-targeting control siRNA (control siRNA that is not complimentary to any human transcripts), and transcript levels were measured by qPCR. Here, T cells were cultured only in Accell siRNA delivery media as it improved siRNA uptake in siRNA delivery experiments as shown in Fig.2b and Fig. 3b. In addition, we tested the effect of anti-CD3 antibody stimulation to evaluate whether TCR signaling influences Accell siRNA–mediated gene expression knockdown (Fig.4a)

**Figure 4.**
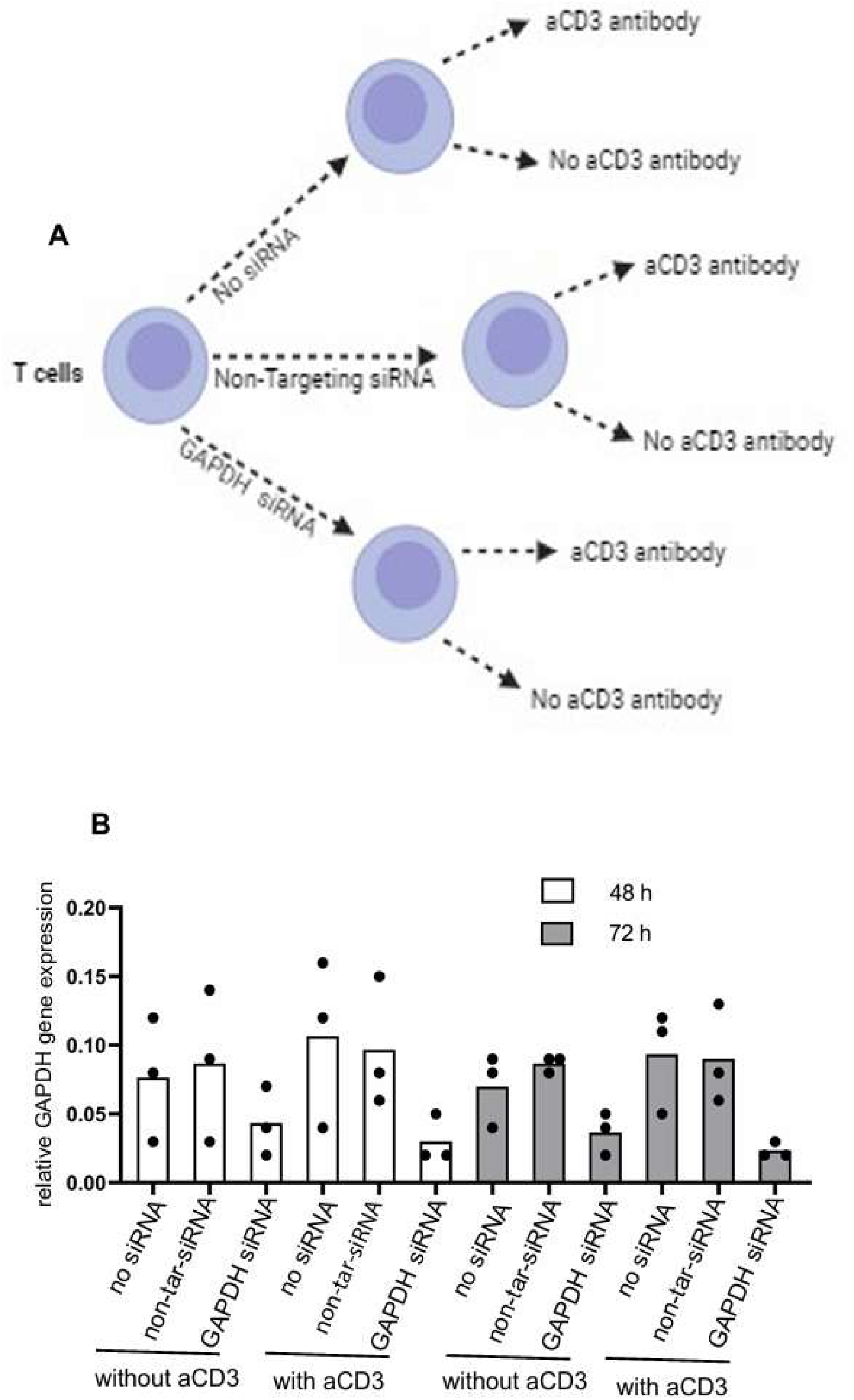
Knockdown of GAPDH in human CD4^+^ T cells using Accell siRNA. (A) Schematic overview of the experimental setup and conditions. (B) qPCR analysis of GAPDH mRNA levels in purified CD4^+^ T cells treated with Accell siRNA targeting GAPDH, in the presence or absence of αCD3 antibody. Cells were cultured in Accell siRNA delivery medium. Non-targeting siRNA was used as a control. Each data point represents an individual donor (n = 3). CD4^+^ T cells were purified from three independent blood donors, treated with siRNA, and analyzed by qPCR. Relative GAPDH expression is shown, normalized to actin as the reference gene.

Accell siRNAs targeting GAPDH achieved robust transcript knockdown. In unstimulated cells, GAPDH expression was reduced by 48% at 48 h and 62% at 72 h (Fig 3b). CD3 stimulation enhanced knockdown efficiency, resulting in reductions of 68% at 48 h and 72% at 72 h (Fig.4b). These data demonstrate that Accell siRNAs can reproducibly knock down endogenous genes in primary CD4+ T cells, with modest augmentation by TCR signaling.

## Discussion

This study establishes a streamlined workflow for Accell siRNA-mediated gene knockdown in freshly isolated, unstimulated human CD4+ T cells. We show that Accell siRNAs are efficiently internalized, maintain high cell viability, and achieve robust transcript silencing. By optimizing delivery conditions, we provide a reproducible method for RNA interference in resting T cells, addressing a major limitation of current transfection-based approaches (Fig.5).

**Figure 5.**
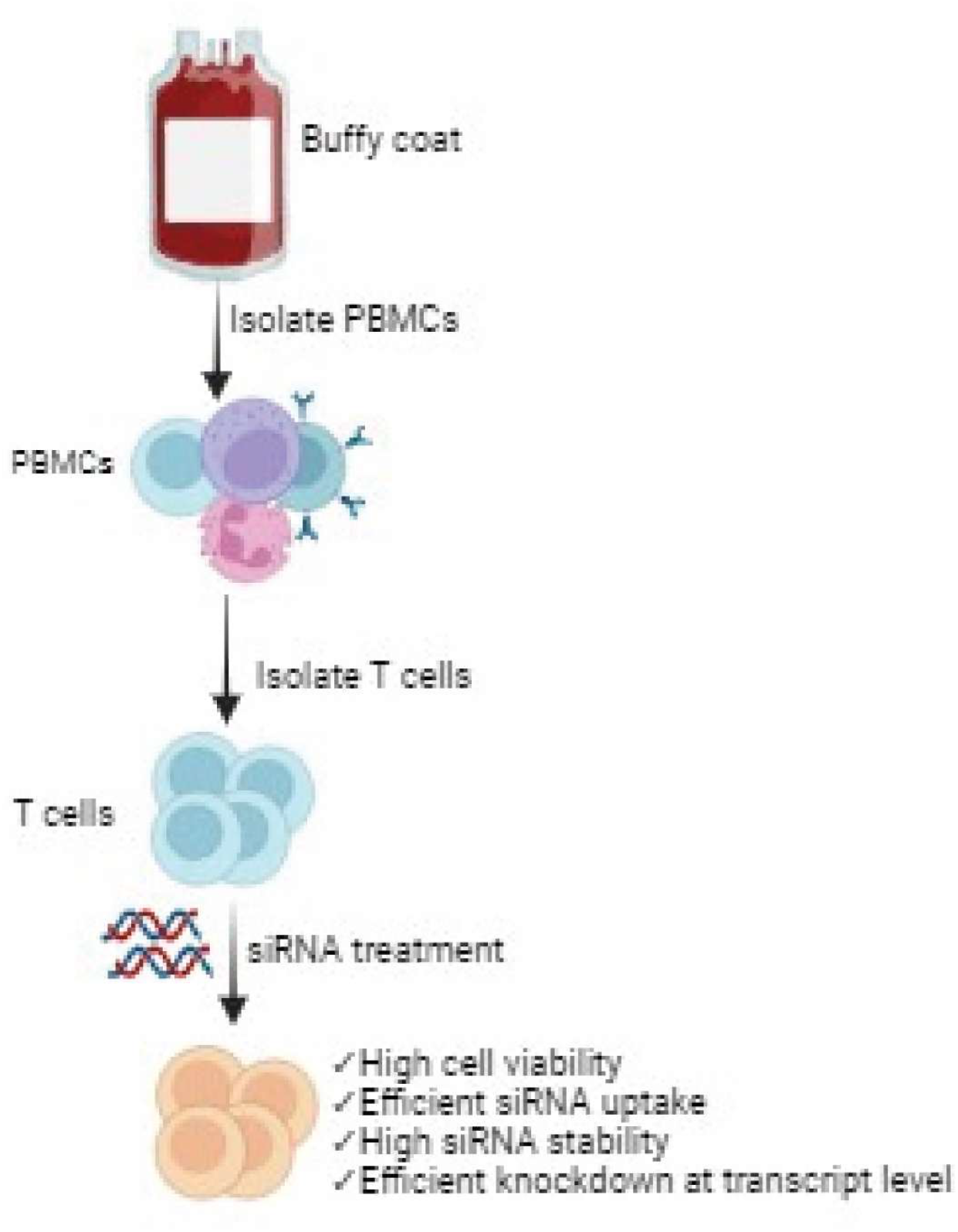
Graphical abstract of the optimized method for Accell siRNA delivery and gene knockdown in primary human CD4^+^ T cells. Created with BioRender

Accell siRNAs have previously been used in a range of hard-to-transfect cell types, including neurons and NK cells, but their application in primary human T cells has remained limited. Prior studies have demonstrated effective delivery in suspension and immune cells such as leukemic models (14), and more recently in complex 3D culture systems (15). Most T-cell–focused applications have relied on expanded or pre-activated T cells (5–13). Our results demonstrate that resting T cells are also amenable to Accell siRNA knockdown, expanding the experimental possibilities for studying T-cell biology under physiologically relevant conditions.

We further highlight culture conditions as an important determinant of siRNA delivery, with Accell delivery medium outperforming standard T-cell media in both magnitude and uniformity of uptake. This provides practical guidance for laboratories planning to adopt Accell siRNAs in primary lymphocytes.

Limitations remain: not all Accell siRNAs yield efficient knockdown, requiring empirical testing of each target. In addition, inducible or rapidly expressed genes may be less suitable, as siRNA uptake and action require sufficient time. Finally, we restricted our analysis to RNA-level silencing; future studies should explore protein-level depletion and functional consequences. Notably, some of these studies have successfully demonstrated protein knockdown in T cells using Accell siRNA-based approaches, underscoring the potential of this platform (5-13).

Despite these caveats, the optimized workflow provides a valuable addition to the experimental toolkit for T-cell research. By enabling efficient gene knockdown without electroporation or viral vectors, Accell siRNAs offer a simple, scalable, and non-disruptive strategy for dissecting gene function in primary human T cells, with implications for studies of autoimmunity, cancer immunotherapy, and T-cell biology.

## Materials and Methods

### Donor material and T-cell isolation

Peripheral blood mononuclear cells (PBMCs) were isolated from buffy coats obtained from healthy donors (Universitetssykehuset Nord-Norge, Tromsø, Norway) under institutional approval. PBMCs were separated by density-gradient centrifugation using Lymphoprep (Stemcell Technologies, #07811). CD4+ T cells were purified by negative selection using a CD4+ T-cell isolation kit (Miltenyi Biotec, #130-096-533) and LS columns with a QuadroMACS separator.

### Cell culture and siRNA treatment

CD4+ T cells (5 × 10^6^) were resuspended in 500 μl of serum-free medium (X-VIVO-15, TexMACS, or Accell delivery medium) and plated in 48-well plates. Cells were treated with 1 μM Accell siRNA (Horizon Discovery) — either a non-targeting control pool (D-001910-10-20) or a GAPDH-targeting pool (D-001930-10-20). Anti-CD3 antibody (1 μg/ml, Tonbo Biosciences #40-0038-U500) was added where indicated. Cells were cultured at 37°C, 5% CO_2_, and harvested at 24–96 h.

### Flow cytometry

For uptake analysis, cells were treated with Accell green-labeled non-targeting siRNA (D-001950-01-20). At 24 h and 48 h, cells were stained with Near-IR Live/Dead dye (Thermo Fisher Scientific) and analyzed on a BD LSRFortessa cytometer. Data were processed in Cytobank, and mean fluorescence intensity (MFI) was used as a measure of siRNA uptake.

### RNA extraction, cDNA synthesis, and qPCR

Total RNA was extracted using the Quick-RNA Miniprep Kit (Zymo Research, #R1055) with on-column DNase I digestion. cDNA was synthesized with the High-Capacity cDNA Reverse Transcription Kit with RNase Inhibitor (Thermo Fisher Scientific, #4374966). qPCR was performed using PowerUp™ SYBR™ Green Master Mix (Thermo Fisher Scientific, #A25918) on a Roche LightCycler 96 instrument. Relative expression was calculated using the ΔΔCT method with Actin as the reference gene (16,17).

### Primer sequences

- GAPDH forward: CTTTTGCGTCGCCAG (140 bp product)
- GAPDH reverse: TTGATGGCAACAATATCCAC
- Actin forward: CTTCGCGGGCGACGAT (103 bp product)
- Actin reverse: CACATAGGAATCCTTCTGACCC

### Step-by-Step Protocol

#### Reagent prep

1. Reconstitute Accell siRNAs to 50 μM in 1× siRNA buffer. Aliquot, store at −20 °C.
2. Pre-warm Accell delivery medium.

#### Cell prep

1. Isolate PBMCs via Lymphoprep gradient.
2. Purify CD4+ T cells using Miltenyi negative selection kit.
3. Count and resuspend 5 × 10^6^ cells in 0.5 mL Accell delivery medium.

#### siRNA treatment

1. Seed cells into 48-well plates.
2. Add siRNA to 1 μM final concentration.
3. Add anti-CD3 (1 μg/mL) if testing stimulation.
4. Incubate 24–96 h at 37°C, 5% CO_2_.

#### Uptake analysis (optional)

- Treat cells with FAM-labeled Accell control siRNA.
- Assess at 24 h and 48 h by flow cytometry with viability staining.

#### mRNA knockdown

- Collect cells at 48 h and 72 h.
- Extract RNA (DNase-treated), synthesize cDNA.
- Perform qPCR for target and reference genes.
- Analyze knockdown using ΔΔCt method.

#### Expected outcomes

- 90–95% viability.
- Strong siRNA uptake by 24 h, sustained at 48 h.
- Robust transcript knockdown by 48–72 h.

#### Notes for siRNA Knockdown Protocol (Accell system)

- Do not add serum or BSA to the Accell siRNA delivery medium, as they inhibit siRNA uptake by the cells.
- Cell culture medium (Accell siRNA delivery medium) can be replaced with serum-supplemented medium after 48 h of siRNA treatment if required.
- siRNA quality can be checked on a NanoDrop to confirm that the vial contains the expected concentration.
- Recommended collection time points:
  - mRNA knockdown: 48 h and 72
  - Note: Optimal time points may vary depending on the target gene. In our experiments, efficient transcript knockdown was observed at both 48 h and 72 h.
- We recommend using siRNA SMARTpools (a mixture of 4 siRNAs targeting the same gene) rather than individual siRNAs for more robust knockdown.

## Acknowledgements

I would like to thank Professor Ole Morten Seternes from the Department of Pharmacy for providing the laboratory space necessary for this research. I am also grateful to the Norwegian Cancer Society for their support, which made this study possible. Special thanks to Tor Brinjar Stuge for providing the MACS separator used for T cell isolation. I sincerely thank Dr. Ali Raja for his assistance with GraphPad Prism in creating the figures for this article. I am also grateful to Dr. Sukumar Namineni for reviewing the manuscript and providing valuable comments.

